# Ex vivo human tumor slices more accurately predict patient responses to an oncolytic virus than in vivo mouse models

**DOI:** 10.64898/2026.09.01.748437

**Authors:** Jose Maldonado, Thomas Wong, Jia-Shiun Leu, Nourhan Abdelfattah, Han Nhat Tran, David Baskin, Robert Rostomily, Nestor Esnaola, Kristin Huntoon, Betty YS Kim, Wen Jiang, Syed Muddassir, Amanda Anderson, Preeti Marie, Alaa Tamim, Natalie Fowlkes Wall, Taku Yoshida, Masamichi Mori, Serguei Soukharev, Shinsuke Nakao, David Menter, Scott Kopetz, Kyuson Yun

**Affiliations:** Department of Neurology, Houston Methodist Research Institute; Naresh K. Vashisht College of Medicine, Texas A&M Health Science Center, Texas A&M University; Department of Neurosurgery, Houston Methodist Research Institute; Department of Surgery, Houston Methodist Academic Medicine; Department of Neurosurgery, The University of Texas MD Anderson Cancer Center; Department of Neurosurgery, University of Arizona and at Southern Arizona VA Health Care System; Department of Radiation Oncology, The University of Texas MD Anderson Cancer Center; Department of GI Oncology, The University of Texas MD Anderson Cancer Center; Department of Veterinary Medicine and Surgery, The University of Texas MD Anderson Cancer Center; Astellas Pharma Inc; Department of Neurology, Weill Cornell Medical College

## Abstract

Immunotherapies, including oncolytic viruses (OV), are promising therapies that can enhance anti-tumor immune responses. However, preclinical success of immunotherapies in mouse models has not always translated to clinical benefit in cancer patients. This study compared preclinical efficacy and mechanism of action for ASP9801, a vaccinia virus expressing IL-7 and IL-12, using mouse models of colorectal cancer (CRC) in vivo and in human organotypic tumor slice models ex vivo. The murine surrogate for ASP9801 significantly reduced tumor volumes in treated and abscopal tumors in two different CRC models in vivo (MC38 and RO100). Treatment efficacy was accentuated when combined with anti-PD1 treatment, and single-cell RNA sequencing analysis revealed depletion of tumor cells and increased T cell infiltration and activation in both treated and abscopal tumors. However, human tissue analysis ex vivo (E-slices) using PDX models and patient samples showed that ASP9801 is not effective in CRC, consistent with clinical trial results. On the other hand, ASP9801 was highly effective in GBM, indicating indication-specific efficacy of ASP9801, and how E-slice assays can be used to identify treatment-sensitive indications. This study demonstrates the superiority of E-slices over mouse models for predicting clinical response and its utility in planning clinical trials.

## Introduction

Immunotherapies, including immune checkpoint inhibitors and chimeric antigen receptor (CAR) - T cell therapy, can provide remarkable clinical benefit to patients with high tumor mutational burden across different cancer types^1–3^. However, the response rate varies widely by cancer type, and remains low overall^4^, and 60-90% of immunotherapies fail in clinical trials despite promising results from preclinical studies. There are multiple reasons for this failure, including the importance of the local tumor microenvironment (TME) in each patient tumor^5^. The poor clinical predictability of preclinical models, including mouse models, human cell lines, and 3D models that lack the native TME in each patient is a major bottleneck in successful development of new anti-cancer drugs in general. Therefore, a more rigorous preclinical evaluation of new therapies using clinically relevant models with intact TME is needed to improve the success rate of new immunotherapies and other anti-cancer agents.

Multiple efforts to augment the efficacy of immunotherapy by reprogramming the immunosuppressive tumor microenvironment (TME) are underway, including rationally designed combinatorial strategies, cytokine therapy, and oncolytic viruses (OV). OVs can both selectively lyse tumor cells and promote immunogenic cell death to improve the anti-tumor immune response^6–9^. Currently, OVs have been approved for the treatment of melanoma, glioblastoma, and nasopharyngeal carcinoma^6,10,11^. OVs have the potential to overcome the immunosuppressive TME by promoting immunogenic cell death, leading to enhanced neoantigen release, and T cell infiltration and activation. OVs can be further engineered to deliver multiple eukaryotic transgene payloads to support local immune cell activation in the TME while avoiding systemic toxicities reported with traditional cytokine therapy^6,7,12–14^. We previously reported an oncolytic vaccinia virus, ASP9801, expressing both interleukin-7 (IL-7) and interleukin-12 (IL-12) to overcome systemic toxicity and to reprogram the TME^12,13^. Introduction of *Il-7* and *Il-12-*expressing ASP9801 surrogate elicited a measurable response in CRC tumors in vivo that did not respond to either of the immune checkpoint inhibitors αPD-1 or αCTLA-4^13^. These in vivo results in mouse models suggested that ASP9801 reprograms the TME to promote T cell infiltration^12,13^ and supported its clinical development.

Here, we compared the preclinical efficacy of ASP9801 in mouse models of colorectal cancer (CRC) and in human ex vivo models, using an organotypic tumor slice culture method (E-slice assays) from patient derived xenografts (PDX) and fresh human tumors. The ASP9801 surrogate was effective in reducing treated and abscopal tumor volumes in two mouse models of CRC, with increase anti-tumor immune response as intended. However, ASP9801 had limited efficacy in human tumors ex vivo, except in GBM, consistent with the recently published clinical trial results for ASP9801^15^. This study demonstrates that clinically relevant ex vivo human cancer models are more predictive of clinical responses than in vivo mouse models and shows that E-slices can be used to select tumor types in which the new treatment may be efficacious.

## RESULTS

### Comparison of sensitivity to ASP9801 in human and mouse cell lines

To determine the efficacy of ASP9801 in different cancer types, we assessed the sensitivity of various human cancer cell types to viral infection by measuring changes in cell viability in vitro (**Figure 1A).** Established human cancer cell lines (Detroit 562 [head and neck: H&N], A549 [lung], HCT116 [colorectal], and U87 [GBM]) and previously characterized mouse colorectal cancer cell lines (CT26 and MC38) were infected with control or species-matching ASP9801 virus **(Figure 1A, B, Supplementary Table 1)**. Compared to untreated controls, viral transduction of both control and ASP9801 virus induced a statistically significant reduction in viability in all cell lines at both MOIs tested. In the absence of immune cells, ASP9801 treatment showed no statistically significant difference compared to control virus for any of the cell lines tested as anticipated.

**Fig. 1.**
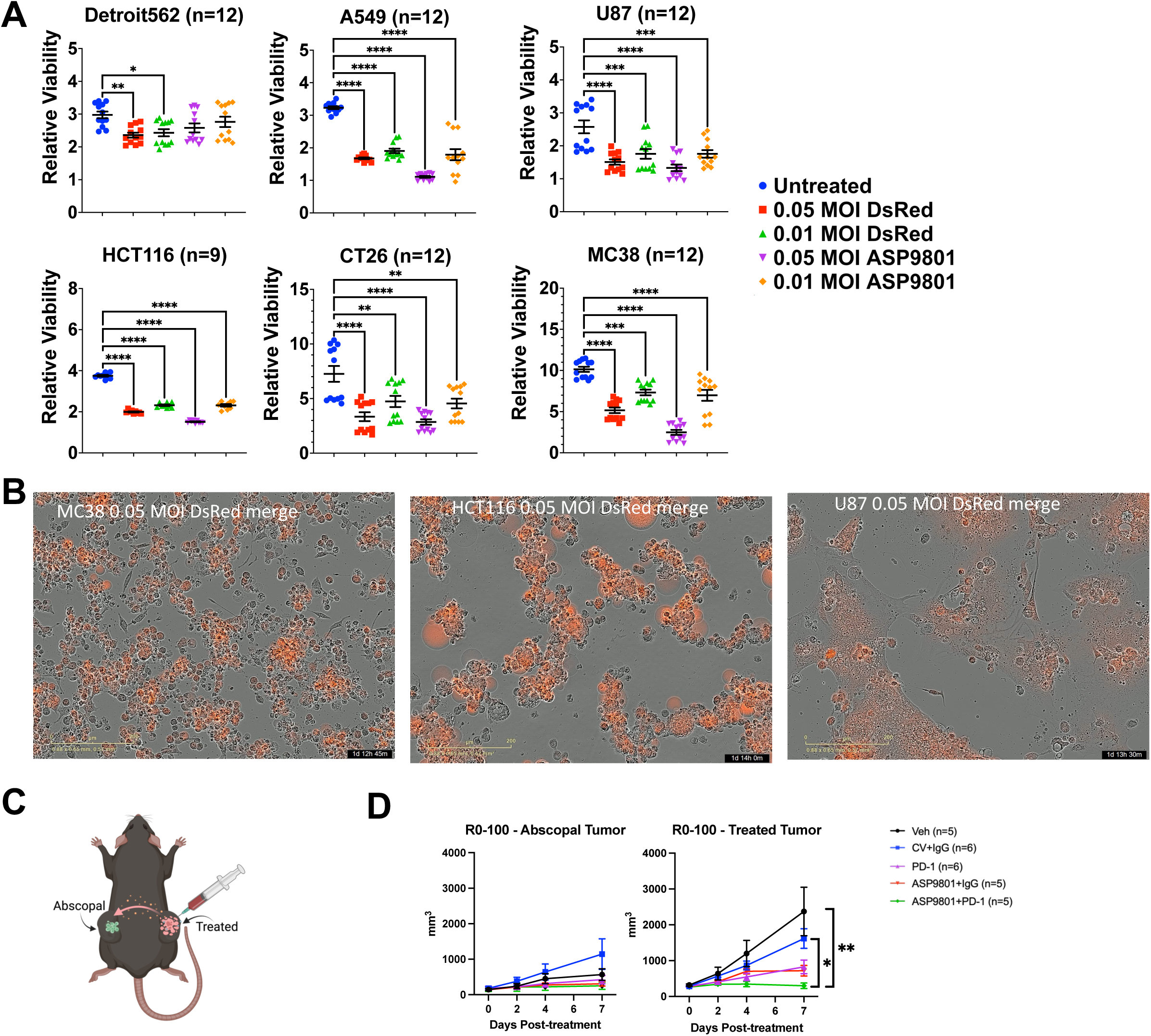
Human and mouse cancer cell lines treated with ASP9801 in vitro and in vivo. A) Changes in viability (relative to t=0) upon control (DsRed) or ASP9801 virus treatment, measured 48-hours after virus exposure in culture. n=12 technical replicates, split across two independent experiments. B) Merged brightfield and fluorescence Incucyte images of cell lines treated with control (DsRed) virus at 36 hours after virus exposure. Scalebar: 200 µm. C) A schematic of *in vivo* ASP9801 treatment study design in bilateral flank tumor mouse model. D) Tumor size changes of RO-100 tumors after treatment with Vehicle control, CV + Iso, aPD-1, ASP9801 + Iso, and ASP9801+aPD-1 to day 7 post treatment, when most control group mice reached humane endpoint and had to be euthanized. Error bars represent SEM. *p*-values determined by one-way ANOVA with Tukey’s correction for multiple comparisons. *p < 0.05, **p < 0.01, ***p < 0.001, ****p < 0.0001. n=5-6 biological replicates per group.

### Intratumor ASP9801 treatment synergizes with *a*PD-1 therapy

To test ASP9801 efficacy in in vivo models with an intact immune system and to evaluate whether it can synergize with αPD-1 treatment, we compared control vs ASP9801 virus treatment in two syngeneic murine CRC models in the C57BL6/J (B6) genetic background: MC38 and RO100. MC38 is a well-characterized mouse CRC model in the B6 background, while RO100 is a newly characterized murine organoid model, also in B6 background, generated from a spontaneous tumor in *APC* and *TP53* mutant mice (Scott Kopetz’s Lab) that recapitulates the common oncogenic events in human CRC.

To study ASP9801 in these CRC models, we implanted either MC38 cells (10^6^ per side) or RO-100 cells (10^7^ per side) bilaterally in B6 mice to develop allografts on the left and right flanks (**Figure 1C**). When tumors on both sides reached approximately 250-300 mm^3^ in size, we treated the right flank tumor with either vehicle, control virus (CV) + IgG, αPD- 1 alone, ASP9801 + IgG, or ASP9801 + αPD-1. ASP9801 or CV virus were injected once with 2 x 10^7^ PFU in 30 μL PBS, and αPD-1 or IgG control antibodies (200 μg) were administered twice/week *via* intraperitoneal injection (IP). Both virus-treated and contralateral tumors were monitored for tumor growth, and both tumors and blood were harvested at day 7 post treatment to determine the treatment-induced changes in the TME. In RO100-injected mice, tumor growth was reduced after treatment with either ASP9801 + IgG or ASP9801 + αPD-1 on treated and abscopal sides (**Figure 1D**). A similar trend was observed in the MC38 model although the tumor size reduction did not reach statistical significance 7 days after treatment (**data not shown**).

### ASP9801 treatment increases systemic and local anti-tumor immune response

To assess the immune modulatory effects of ASP9801, we performed high parameter spectral flow cytometry analyses on tumor digests and PBMCs (**Figures 2-3, Supplementary Figure 1 and 2**). In the virus-treated tumor, OV treatment increased CD3^+^ and CD8^+^ T cell infiltration (**Figure 2A, B**), with the largest and statistically significant increase in the ASP9801+αPD-1 combination treated group. On the other hand, the proportion of PD-1^+^ CD8 T cells was significantly decreased in the combo treated group, from nearly 80% of T cells in the vehicle group to around 20% in ASP+αPD- 1 group (**Figure 2C**). Conversely, the proportion of CD69^+^ CD8 T cells significantly increased in the ASP9801+αPD-1 group (**Figure 2D**) while CD69^+^PD-1^+^ CD8 T cells decreased in the combination treated group (**Figure 2E**). CD69^+^PD-1^-^ population increases ∼3 fold in the combination treated group compared to all other groups (**Figure 2F**), suggesting that the combination treatment increased tumor reactive CD8 T cells with less exhausted phenotype. CD4^+^ T cells constitute less than 4% in the vehicle treated group (**Figure 2G**), and there was a trend towards increased frequency in the virus treated groups, although not statistically significant (**Figure 2G**). However, the fraction of CD25^+^PD-1^+^ CD4 T cells (Tregs) significantly dropped from ∼40% in vehicle treated to less than 10% in the combination treated group (**Figure 2H**). Together, these results suggest that oncolytic virus treatment of MC38 tumors increases cytotoxic CD8 T cell infiltration, especially when combined with αPD-1 treatment, and the combination treatment significantly decreased PD-1^+^ exhausted CD8 T cells and Tregs, compared to vehicle control treated mice.

**Figure 2.**
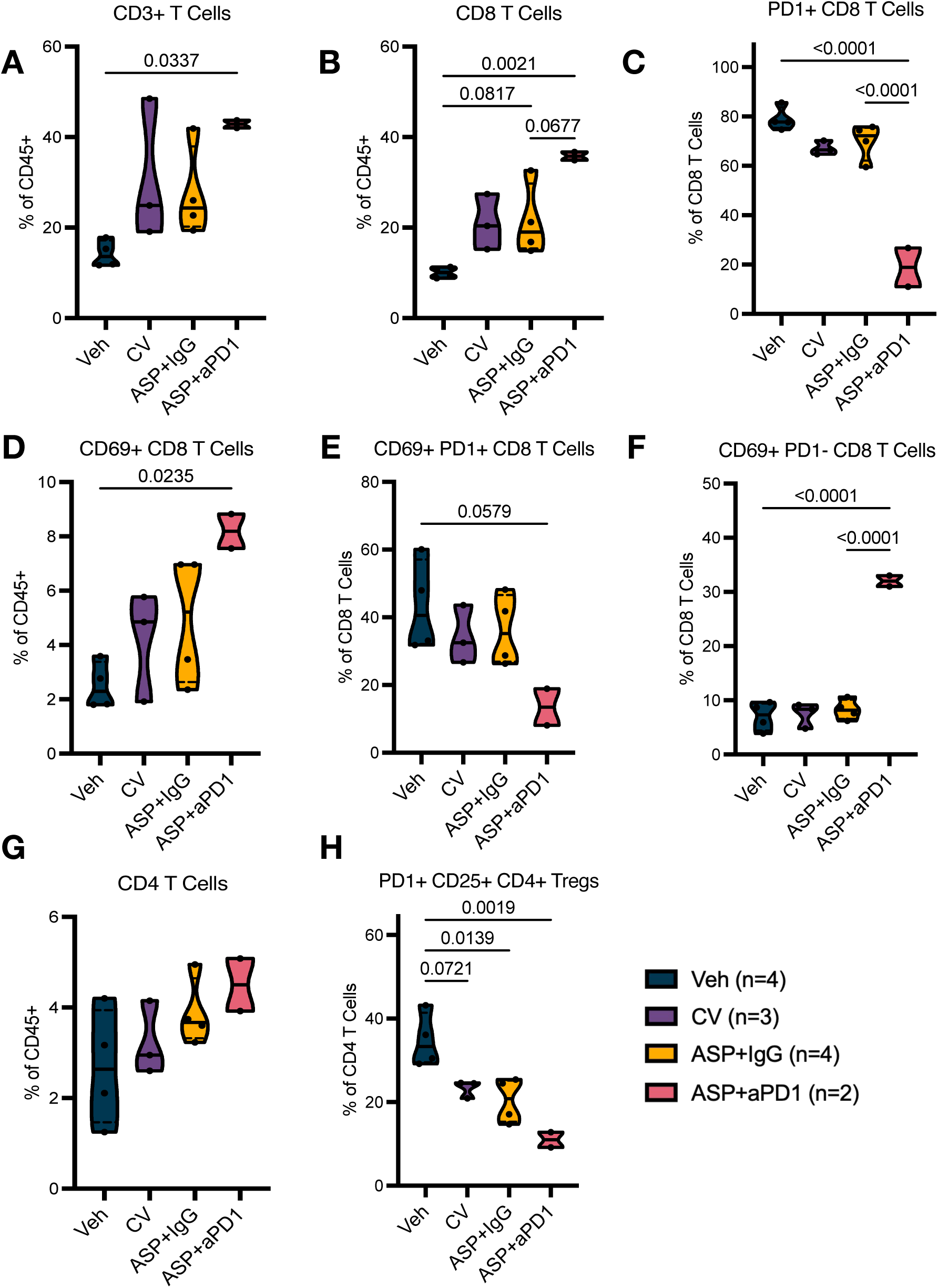
Intratumor ASP9801 treatment synergizes with *a*PD1 therapy to increase T cell infiltration/activation. **A-I**, Virus treated MC38 murine CRC tumors were analyzed by spectral flow cytometry with: CD3^+^ T cells (A) and CD8^+^ T cells (B) as a fraction of CD45^+^ immune cells, (C) PD1^+^ CD8 T cells as a fraction of CD3^+^ T cells (D) CD69^+^CD8^+^ T cells as a fraction of CD45^+^ cells (E), CD69^+^PD1^+^ CD8 T cells and (F) CD69^+^PD1^-^ CD8 T cells as a fraction of CD8 T cells. (G) CD4^+^ T cells as a fraction of CD45^+^ cells (H) PD1^+^ CD25^+^ CD4 Treg cells as a fraction of CD4 T cells, at 7 days post treatment. n=2-4 per treatment group, indicated by dots. *p*-values determined by one-way ANOVA with Šidák’s correction for multiple comparisons. For violin plots, ends represent minimum and maximum values, solid line represents median value, and dashed lines represents quartiles. n=2-4 biological replicates/group.

**Figure 3.**
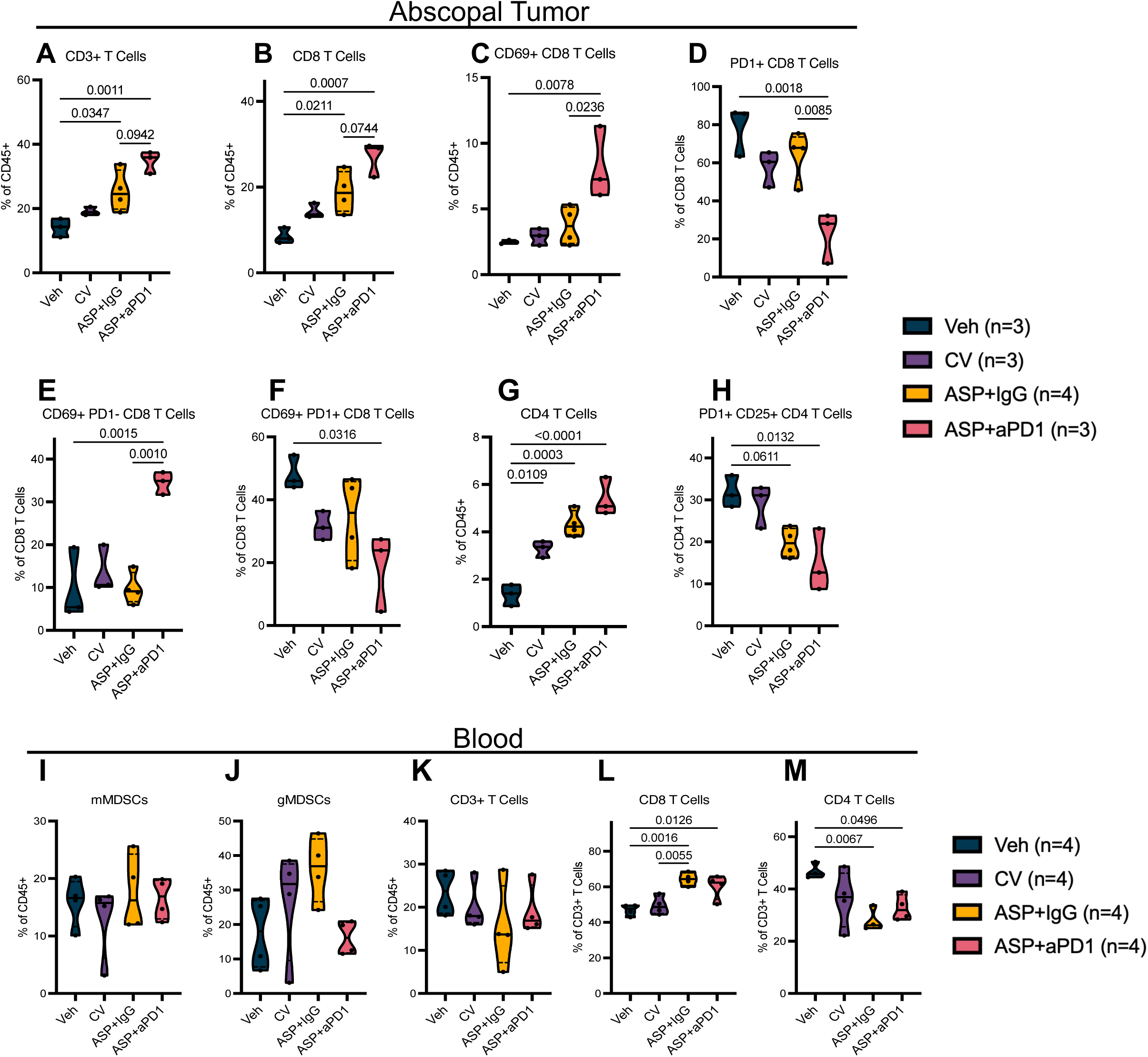
ASP9801 treatment promotes an abscopal immune response that synergizes with *a*PD1. **A-H**, Untreated MC38 CRC tumors were analyzed by spectral flow cytometry at the same time as treated tumors presented in Figure 2. (A) CD3^+^ T cells and (B) CD8^+^ T cells as a fraction of CD45^+^ immune cells. (C) PD1^+^ CD8 T cells as a fraction of CD8 T cells (D) CD69^+^ CD8 T cells as a fraction of CD45^+^ cells, (E) CD69^+^PD1^+^ CD8 T cells, (F) CD69^+^PD1^-^ CD8 T cells, (G) CD4^+^ T cells, and (H) PD1^+^CD25^+^ CD4 T cells from abscopal MC38 tumors at 7 days post treatment. n=3-4 biological replicates/group. **I-M**, Flow cytometry analysis of circulating (I) mMDSCs, (J) gMDSCs, (K) CD3^+^ T cells, (L) CD8^+^ T cells, and (M) CD4^+^ T cells in the blood of the same mice at 7 days post treatment. *p*-values determined by one-way ANOVA with Šidák’s correction for multiple comparisons. For violin plots, ends represent minimum and maximum values, solid line represents median value, and dashed lines represents quartiles. n=4 biological replicates/group.

A key advantage of OV therapy is the activation of tumor antigen-specific T cells (stemming from the treated side) that can have anti-tumor activity in distant (abscopal) tumors. Hence, we analyzed immune infiltrates in abscopal tumors at the same time point as the virus-treated tumors. ASP9801 treatment had analogous effects in abscopal tumors as in the virus-treated tumors: significantly increased CD3, CD8, and CD4 T cells in ASP9801 and combo treated groups, compared to vehicle control (**Figure 3A, B, G**). Like treated tumors, the proportion of CD69^+^ CD8 T cells increased (**Figure 3C**) and PD- 1^+^ CD8 T cells decreased significantly in the combo treated group compared to both vehicle control and ASP9801 treatment alone (**Figure 3D**). Activated CD69^+^ CD8 T cells increased significantly in the combo group, primarily due to increased CD69^+^PD-1^-^ CD8 T cells (**Figure 3E**) while CD69^+^PD-1^+^ CD8 T cells were reduced in the OV treated groups (**Figure 3F**). In the CD4 T cell compartment, CD4+ T cell frequency increased significantly in ASP9801 and the combo group (**Figure 3G**) while the proportion of Tregs decreased significantly with the combo treatment (p=0.0132) but did not quite reach significance in the ASP9801 alone group (p=0.0611, **Figure 3H**).

In contrast to T cells, myeloid cell changes were less significant. Antigen presenting IAIE^+^/MHC-II^+^ myeloid cells as a proportion of all myeloid cells were significantly increased in the combo group compared to the ASP9801 group in the abscopal tumor but did not reach significance in the treated tumor (p=0.0925, **Supplementary Figure 2A, B**). The frequencies of IAIE^-^Ly6g^-^Ly6c^high^ monocytic myeloid derived suppressor cells (mMDCSs) were reduced in both the treated and abscopal tumors, but they did not reach statistical significance **(Supplementary Figure 2A, B)**.

We assessed circulating immune cells as well and found no significant changes in the proportions of mMDSCs, gMDSCs (IAIE^-^Ly6g^high^Ly6c^int^), or CD3^+^ T cells, as a proportion of all CD45^+^ cells (**Figure 3I-K**). However, the CD8 T cell fraction in circulation was increased in the ASP9801 and combo treated mice while the CD4 T cell fraction was significantly decreased (**Figure 3L, M**). Together, these results indicate that intratumor ASP9801 treatment induces a strong anti-tumor immune response in treated and untreated lesions by increasing T cell infiltration and activation, which synergizes with systemic αPD-1 treatment.

### Anti-tumor immune response is enhanced by ASP9801 + αPD-1 combination

To gain insights into treatment-induced changes at the molecular level, we performed single-cell RNA sequencing (scRNAseq) analysis on MC38 tumors from both treated and abscopal sides, 7 days post-treatment. We analyzed 40,871 aggregated single cells that passed quality control from 8 treatment groups (pooled from 3 mice for each treatment group). Unsupervised clustering of all aggregated cells revealed 27 clusters with distinct gene expression patterns **(Figure 4A).** We used a combination of top differentially expressed (DE) genes, SingleR automated cell type annotation, and CNV prediction to identify 13 broad cell types (**Figure 4A, Supplementary Figure 3A-C**), including Neutrophils (*S100a8*, *S100a9*, *Hdc*), B cells (*Cd79a*), T cells (*Cd3e*), cDC1 (*H2-Eb1*, *Clec9a*), cDC2 (*H2-Eb2*, *Fscn1*, *Ccr7*), pDCs (*Siglech*, *Klk1*), Macrophages (*Itgam*, *Apoe*, *C1qa*), Monocytes (*Plac8*, *Vcan*, *Chil3*), Endothelial cells (*Vwf*, *Esam*, *Cdh5*), Cancer-Associated Fibroblasts (CAFs) (low CNV-*Col1a2*, *Bgn*), and Tumor clusters (high CNV, MC38 markers).

**Figure 4.**
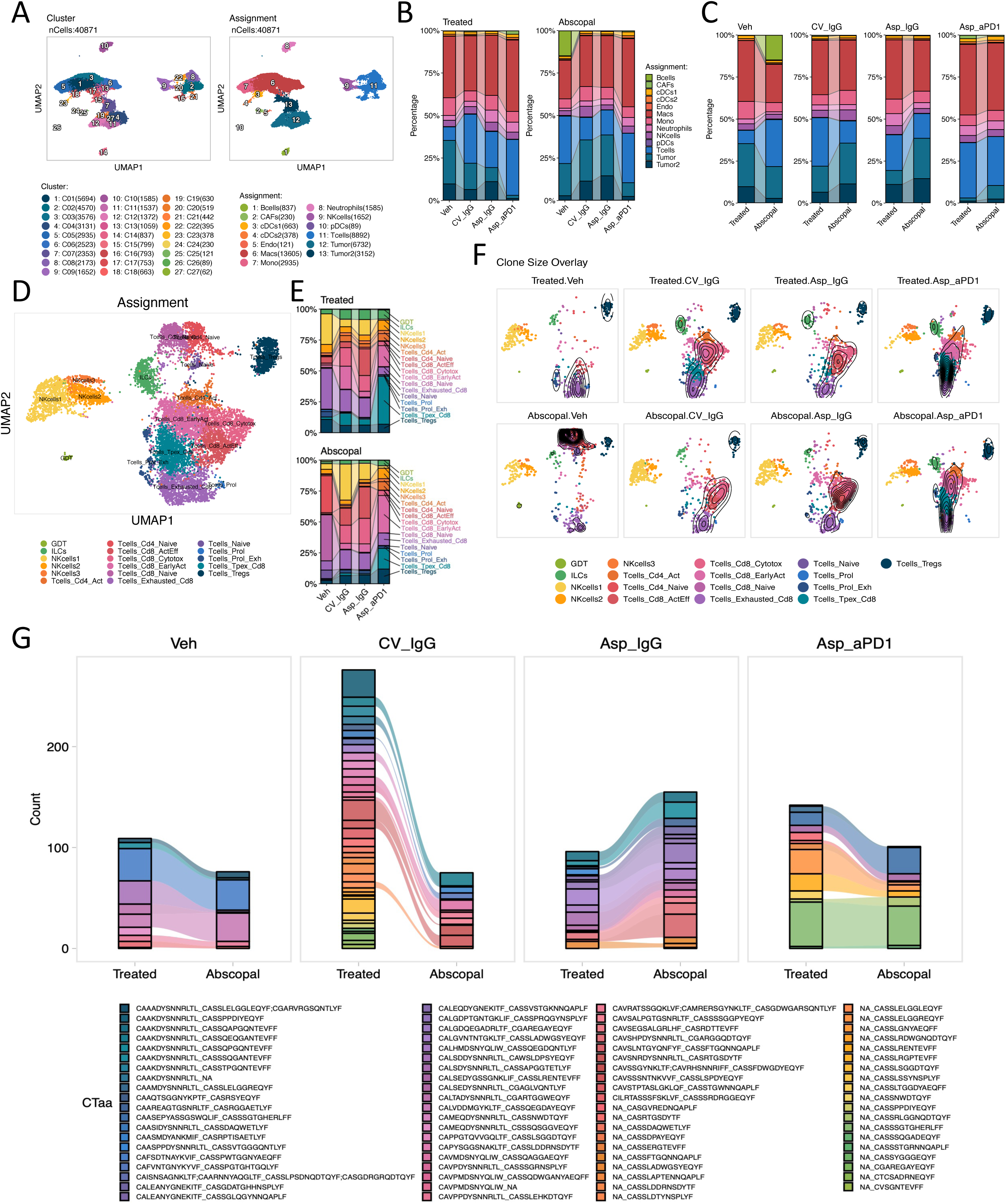
Single-cell landscape of MC38 primary and abscopal tumors. **A,** UMAP projection of 40,871 integrated single cells, colored by cluster (left) and cell type annotation (right). n=3-4 pooled biological replicates/group. **B,** Stacked bar plots displaying cell-type shifts across treatment groups (Veh, CV_IgG, Asp_IgG, Asp_aPD1), stratified by tumor site (Treated vs. Abscopal). **C,** Data from panel B re-visualized in a pairwise manner per treatment group. **D,** De novo clustering and UMAP projection of T and NK cells (n= 10,495) colored by cell type. **E,** Stacked bar and trend plots displaying T and NK cells cell-type shifts across treatment groups. Color coding same as in D. **F,** TCR clone size density mapping on T-cell UMAPs identifying regions of significant clonal expansion. **G,** Alluvial plot depicting the distribution and sharing of TCR clones across experimental groups filtered by clones common between at least 3 samples.

Consistent with our tumor volume measurements, normalized stacked bar graphs show a marked reduction in the tumor cell fraction in OV treated groups (**Figure 4B,C**). To further analyze treatment-induced changes in T and NK cell states, we performed de novo clustering of T cells and NK cells (**Figure 4D, Supplementary Figure 4A,B**). While all viral treatments increased the percentage of infiltrating CD8 T cells, the molecular profile of these T cells was distinct by treatment group. CD8 T cells in ASP9801-treated and abscopal tumors were more enriched with cytotoxic and activated CD8 T cells, compared to a relative enrichment of early activated CD8 T cells in CV-treated tumors (**Figure 4E**). Notably, while the vehicle treated tumors were enriched with terminally exhausted CD8 T cells, ASP9801 + αPD-1 combination tumors had increased precursor exhausted CD8 T cells in both the treated and abscopal tumors (**Figure 4E,F**). Clonotype analysis through TCR sequencing showed that OV treatment increased the number of clones (**Figure 4F,G, Supplementary Figure 5A-B**). The largest clones in each treatment group were unique and not shared across different treatment groups, both in the treated and abscopal sides (**Figure 4G, Supplementary Figure 5A-B**). When treated and abscopal tumors are compared pairwise by treatment group, most of the large clones were shared between the treated and abscopal tumors (**Figure 4G, Supplementary Figure 5**).

### ASP9801 oncolytic reduces viability in patient-derived xenograft (PDX) slices

In light of the in vivo mouse tumor model results demonstrating anticipated anti-tumor immune response, we assessed the potential clinical efficacy of ASP9801 in human tumors using a novel 3D tumor slice culture method called E-slices^16–18^. The E-slice method maintains an intact TME ex vivo and has been shown to provide a more accurate prediction of treatment response to chemotherapy plus pembrolizumab in triple-negative breast cancer (TNBC) patients^16–18^. We first tested in PDX models generated by injecting patient-derived cells of either CRC, H&N cancer, or GBM into the flanks or brains of immune-deficient NSG mice **(Figure 5A, Supplementary Table 2).** PDX derives E-slices were infected with control (DsRed) or ASP9801 with 10,000 PFU and viability changes at day 8 and day 12 were measured (**Figure 5B-D**). Unlike mouse models, only two of eleven CRC PDX models (C1141, and B8101) responded to ASP9801 treatment with significantly reduced viability (**Figure 5B**). In addition, one of three H&N PDX models and one GBM PDX model showed significant reduction in viability upon either virus treatment (**Figure 5C,D**).

**Fig. 5.**
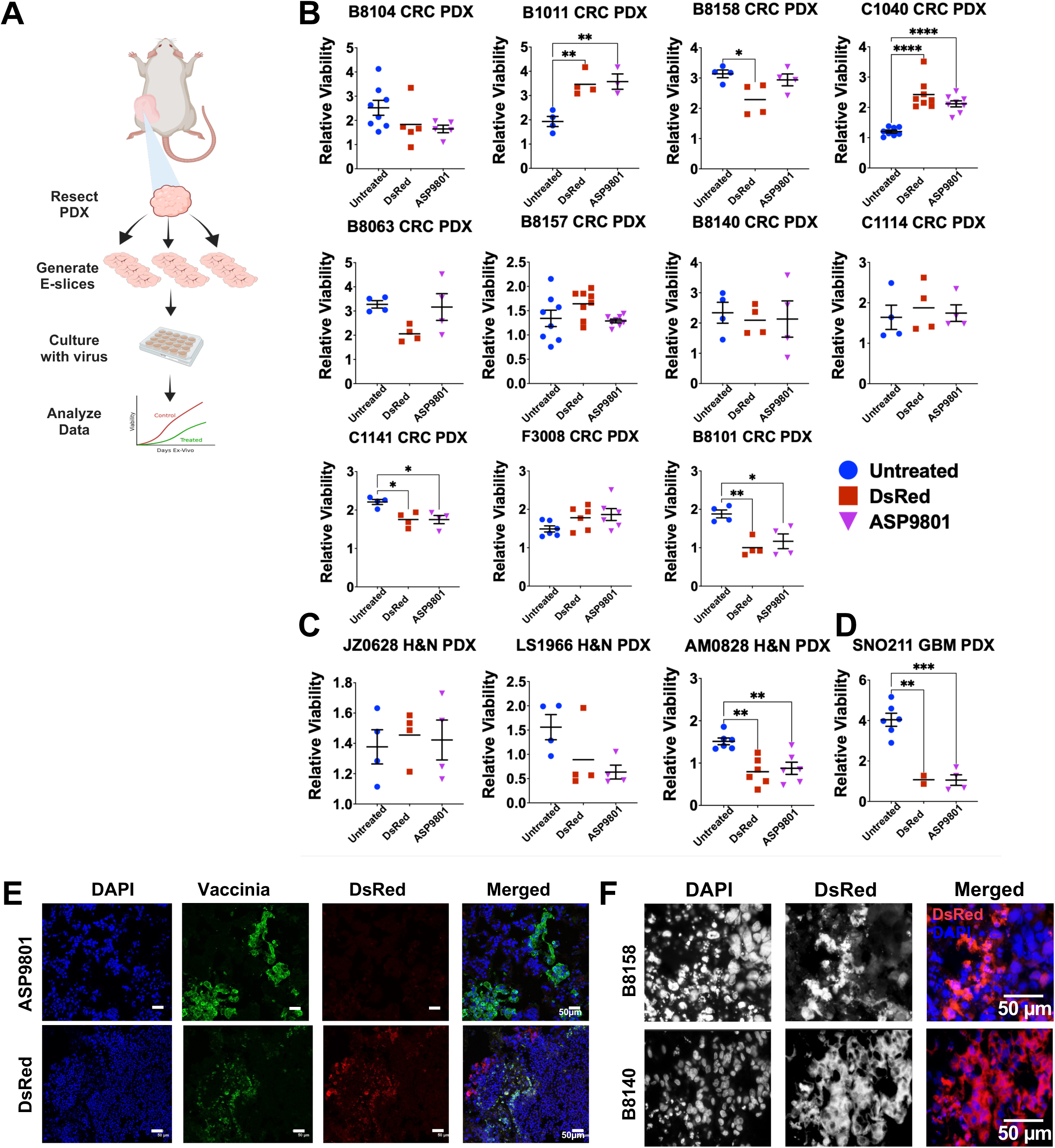
Patient Derived Xenograft E-slice response to ASP9801. **A-D**, (A) Experimental Schema. (B-D) Relative viability change (normalized to baseline) in untreated, control virus (DsRed), or ASP9801 treated at day 8 (presented for B8157, C1040) or day 12 (remaining samples) *ex vivo* (DEX). (B) colorectal cancer PDX models, (C) head and neck PDX models, and (D) glioblastoma PDX models. Each model was tested 2-3 times from different PDX host mice and in duplicates or triplicates in each experiment, depending on available tissue. p-values determined by one-way ANOVA with Tukey’s correction for multiple comparisons, error bars represent SEM. *: p < 0.05, **: p < 0.01, ***: p < 0.001, ****: p < 0.0001. **E,** Immunofluorescence (IF) detection of vaccinia virus in a CRC PDX model at 8 DEX. Scalebar: 50 µm. **F,** IF analysis of DsRed control virus infected cells in a responder (B8158) and non-responder (B8140) CRC PDX models showing differences in pyknotic nuclei and dying cells in responders. Scalebar: 50 µm.

Immunofluorescence staining of E-slices showed presence of vaccinia virus infected cells (**Figure 5E**) and most infected cells were dying/dead in B8158 (sensitive model) but not in B8140 (resistant model, **Figure 5F**), suggesting that oncolytic ability of vaccinia virus may vary in different patient tumors. Notably, most human CRC PDX models did not show significant reduction in viability, unlike mouse models. However, since these PDX models were generated in immune-deficient NSG hosts and lacked immune cells that could respond to IL7 or IL12 from ASP9801, we next tested E-slices derived from fresh patient samples with intact TME to assess whether the presence of immune cells enhances the efficacy of ASP9801.

### Effects of ASP9801 on patient tumor slices

To evaluate ASP9801 efficacy in a more clinically relevant model, we tested fresh patient tumor samples with intact TME by generating E-slices from fresh surgical discard material from CRC, GBM, Astrocytoma (AC), and breast cancer brain metastasis (BCBM) **(Figure 6A-D, Supplementary Table 3).** E-slices were infected with 10,000 PFU of control DsRed virus or ASP9801. In the 4 CRC samples tested, ASP9801 treatment induced little to no significant reduction in viability, consistent with our PDX models results (**Figure 6A**). In contrast, 4 of 5 GBM patient samples, and 1 BCBM showed reduced viability to either or both viral treatments (**Figure 6B,D**), although one GBM sample did not reach statistical significance due to limited tissue availability. These results suggest that vaccinia OVs may have higher efficacy in GBM.

**Fig. 6.**
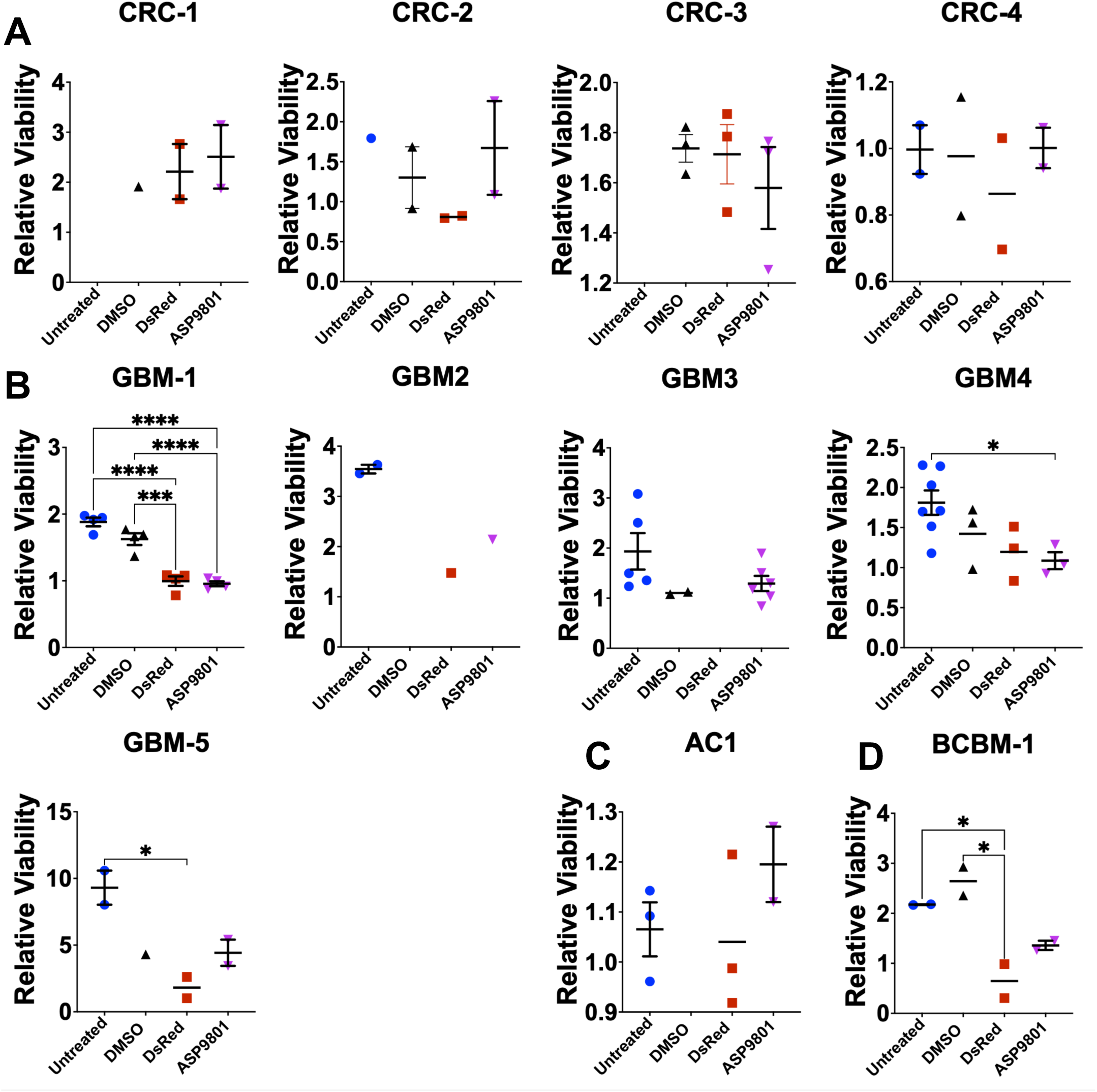
Patient tissue and heterogeneous mouse GBM model responses. **A-D**, E-slice viability readings for (A) colorectal cancer patient samples, (B) GBM patient samples, (C) Astrocytoma patient sample, and (D) breast cancer brain metastasis sample. 10,000 PFU control virus or ASP9801 was used to treat E-slices. Missing data points are due to the limited size of patient tumor samples. CRC: Colorectal Carcinoma, GBM: Glioblastoma, AC: Astrocytoma, BCBM: Breast Cancer Brain Metastasis. p-values determined by one-way ANOVA with Tukey’s correction for multiple comparisons, error bars represent SEM. *p < 0.05, **p < 0.01, ***p < 0.001, ****p < 0.0001.

### Potential biomarkers of response

Motivated by the significant response in GBM patient tumors, we sought to identify potential biomarkers using conditioned media from patient tissue E-slices. E-slices are cultured in chemically defined, serum-free media. We harvested conditioned media from E-slice experiments and performed a targeted cytokine analysis of conditioned media from responders (showing significant reduction in viability) and non-responders, using the Human Cytokine Array Q640 multiplex immunoassay to monitor 640 cytokine levels **(Figure 7, Supplementary Table 4).** To confirm successful ASP9801 viral infection and cytokine expression, we first compared IL-7, IL-12p40, and IL-12p70 levels **(Figure 7A– C)** across different treatment arms. IL-7, IL-12p40 and IL-12p70 levels were elevated only in ASP9801 treated E-slices in all samples, indicating successful infection, expression and secretion of the transgene products. Additionally, we measured the correlation between treatment response (viability change) and IL-7, IL-12p40, and IL-12p70 levels in the matching samples, and observed a significant and strong positive correlation between secreted IL7 and IL12 levels and changes in viability in each sample **(Figure 7D)**.

**Fig. 7.**
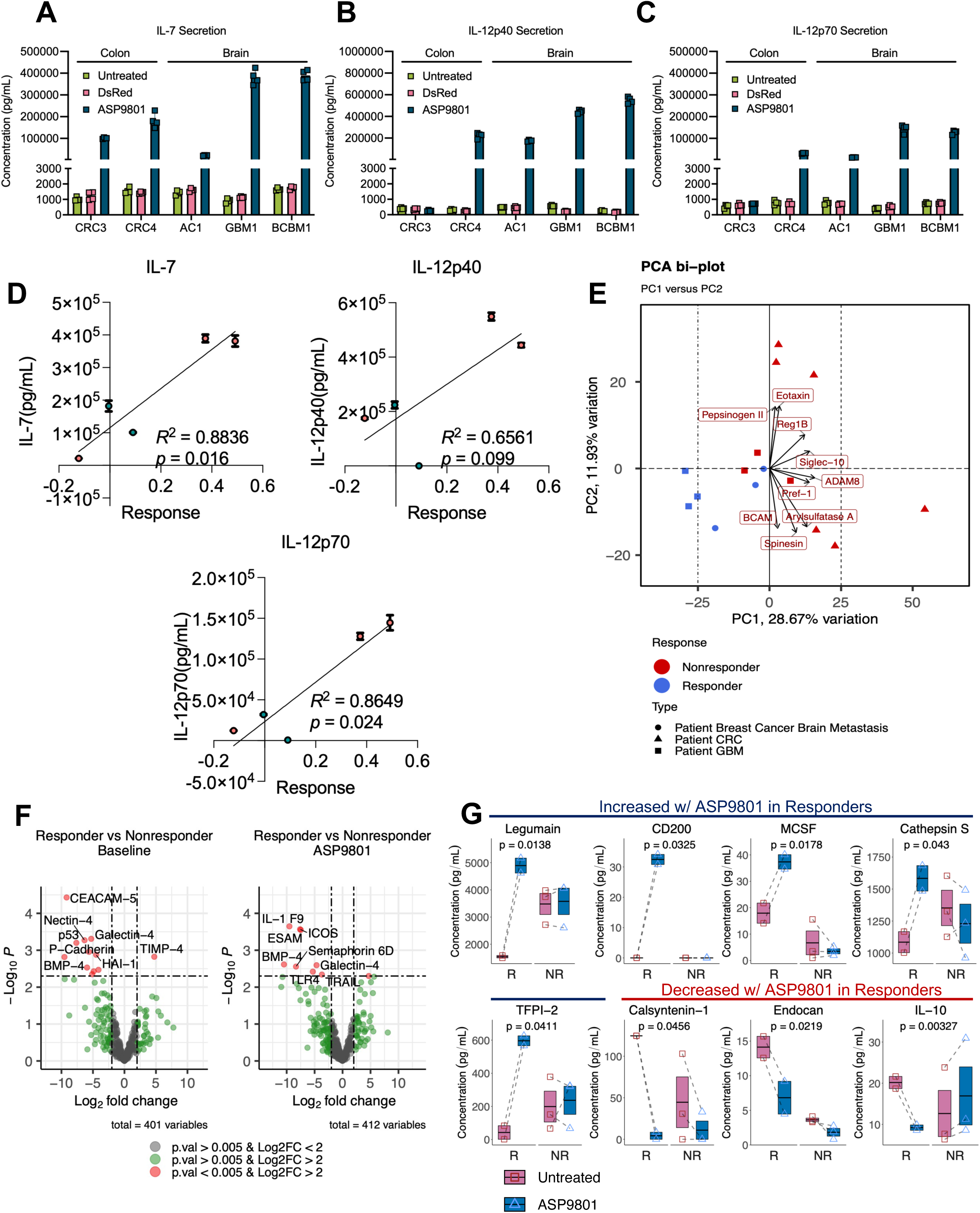
Cytokine Analysis of conditioned media from patient tumor E-slices. **A-C**, Cytokine array quantitation for (A) IL-7, (B) IL-12p40, and (C) IL-12p70 cytokines from tumor slice conditioned media. Technical replicates are plotted here (n=4/treatment). Error bars represent SEM. CRC: Colorectal Carcinoma, GBM: Glioblastoma, AC: Astrocytoma, BCBM: Breast Cancer Brain Metastasis. **D**, Pearson correlation between E-slice response and secreted IL-7, IL-12p40, and IL-12p70 levels in the conditioned medium from ASP9801 treated E-slices. E-slice response was calculated as percent change in viability compared to the untreated sample. Mean for technical replicates (n=4) plotted for each sample; error bar represents SEM. R^2^ and p-values determined from Pearson correlation test. **E,** Principal Component Analysis of patient sample cytokine array data. Top features driving variability are displayed. **F,** Volcano plot showing up- and down-regulated cytokines in responders vs non-responders from conditioned media of baseline/untreated control patient samples (left) and ASP9801 treated patient samples (right). Cytokine array concentrations were log normalized and filtered to exclude low expressing cytokines and bottom quartile variable cytokines. **G,** Paired analysis of secreted cytokine levels comparing change between untreated vs ASP9801-treated samples from each patient, stratified by response status (non-responders (NR), n=3 vs responders (R), n=2). Floating box represents mean with SEM, with horizontal line indicating the mean; individual values are overlaid as points. *p*-values determined by two-sample *t*-test.

To analyze downstream effects of virus-mediated IL-7 and IL-12 expression, we analyzed the levels of other cytokines in the conditioned media, comparing ASP9801-treated to untreated patient tumor slices (**Supplementary Figure 6**). From a principal component analysis (PCA) of cytokine secretion profiles, we observed that responder samples clustered together (**Fig 7E)**. Furthermore, responder tissues at baseline (without treatment) had significantly lower levels of CEACAM-5, BMP4, and Galectin-4, and higher levels of TIMP-4 (**Figure 7F, Supplementary Table 5**). Upon ASP9801 treatment, responder tissues had significantly lower levels of BMP4, ICOS, ESAM, Galectin-4, Semaphorin 6D, and IL-1 F9 (**Figure 7E**). We conducted pairwise comparisons between untreated and ASP9801 treated samples, and discovered that CD200, Cathepsin S, Legumain, MCSF (macrophage colony stimulating factor, also known as CSF-1) and TFPI-2 were significantly elevated in ASP9801 treated E-slice conditioned media compared to baseline-untreated E-slice conditioned media in responders but not non-responders (**Figure 7G**). Conversely, Calsyntenin-1, Endocan, and IL-10 were significantly decreased with ASP9081 treatment in responders, but not in nonresponders (**Figure 7G**). These results suggest that cytokines associated with myeloid cell response and ECM remodeling are associated with response to ASP9801 treatment in E-slices.

## DISCUSSION

Current in vitro models used for predicting clinical efficacy of immunotherapies have significant limitations. There is thus a pervasive need for reliable and efficient in vitro assays that can quickly determine which therapies will be effective for each individual patient. To this end, we propose an ex vivo approach in which tumor slices removed from patients are assayed ex vivo. Using this approach, new anti-cancer therapies, including immunotherapies that require an intact human TME, can be assessed quickly preclinically to measure both in-human efficacy, identify the most promising indications, and gain on- and off-target mechanistic insights and biomarkers for treatment sensitivity and resistance.

Oncolytic viruses (OVs) are commonly viewed today as immune-priming biologics that can kill tumor cells locally, convert “cold” tumors into proinflammatory, T-cell–permissive microenvironments^7^, and are more effective when paired with other immunotherapies rather than used alone. Currently, T-VEC (Imlygic) is an FDA-approved OV for local treatment of unresectable cutaneous/subcutaneous/nodal melanoma^19^. Teserpaturev (Delytact) is conditionally approved in Japan for malignant glioma^20^. Oncorine (H101) has been approved in China to treat nasopharyngeal carcinoma in combination with chemotherapy^21^.

ASP9801 is an oncolytic vaccinia virus designed to treat advanced solid tumors by selectively infecting, replicating in, and lysing cancer cells while sparing healthy tissue. Preclinically, ASP9801 showed very promising results in mouse models, reducing tumor volume in both treated and abscopal tumors, including results from this study. However, human models from this study showed minimal response from CRC and H&N patient samples. On the other hand, promising results were observed in a handful of human GBM samples treated with ASP9801, both in a PDX model as well as in GBM patient samples. These results suggest that ASP901 effectiveness may be cancer type or individual tumor-dependent, highlighting the importance of predictive models to identify in-human efficacy.

Phase 1 clinical trial results from ASP9801 were recently reported by Astellas. ASP9801 showed trends consistent with conversion of a cold tumor microenvironment (TME) to a hot TME, as evidenced by increased tumor-infiltrating lymphocytes (TILs) and elevated PD-L1 expression in post-treatment tumor biopsies^15^. However, they failed to detect clear clinical efficacy^15^, consistent with E-slice measurements in this study. Our findings with patient tumor E-slice assays suggest that GBM is a promising indication for ASP9801; however, GBM was not included in the Phase 1 trial.

Several OVs are currently under clinical investigation for treatment of GBM, and Teserpaturev, an HSV1-based OV, is approved in Japan for patients with malignant gliomas^10,32^. Results from recent studies assessing CAN-3110, an HSV-based OV, in patients with recurrent GBM point to several immune associated factors that may differentiate responders from non-responders to OV treatment, including positive HSV serology pre-treatment and expansion of T cell clonotypes post-treatment, both of which are positively correlated to longer survival^30,33^. Our results from PDX and patient tumor E-slices indicate that GBM responds more favorably to ASP9801 compared to other cancer types. Additionally, preclinical data from others support the rationale for IL-7 and IL-12 to reprogram the TME in treatment of GBM. IL-7 was recently shown to enhance T cell homing to the tumor site in a VLA-4 dependent manner in preclinical mouse GBM models^34^. Similarly, delivery of IL-12 encoded by a replication-incompetent adenoviral vector in mouse and primate GBM models led to improved T cell infiltration and survival^35^. Together, these results support future clinical trials with ASP9801 in GBM.

### Translational outlook

This study highlights the need for human cancer-specific predictive assays, especially for immunotherapy and biologics. Our evaluation of ASP9801 in the E-slice assay shows that this method is valuable for assessing the effectiveness of potential treatment for each patient and possibly for discovering biomarkers of response. Importantly, this study highlights the value of evaluating new therapeutics in clinically relevant human models, particularly to determine efficacy and most appropriate indications for new therapies.

## METHODS

### Human tumor specimen collection

Human tumor tissue was obtained under Institutional Review Board (IRB)-approved protocols (Pro00014547, PA19-0661, & Lab10-0982) at Houston Methodist Hospital, Houston, Texas in accordance with national guidelines. All patients signed informed consent during clinical visits before surgery and sample collection. Patients did not receive compensation in return to their participation in this study. The clinical characteristics of the patient samples are described in **Supplementary Table 3**.

### Mouse models and use

Mice were housed and handled in accordance with the protocols and procedures approved by the HMRI Institutional Animal Care and Usage Committees (protocol # IS00007185, 00001187 and 00001077). Mice were group-housed (up to five animals per cage) in a controlled environment with a standard 12-hour light/12-hour dark cycle. Animals had ad libitum access to food and water. 5–8-week-old C57BL/6J (The Jackson Laboratory, #664) mice were used as hosts for MC38 and RO100 tumor growth. Flank tumors were generated using standard protocol. Briefly, 10^6^ MC38 or 10^7^ RO100 mouse CRC cells were resuspended in 100 µl of Matrigel solution (VWR, # 47743-720) and injected into each side per mouse. Injected mice were monitored daily for tumor growth.

Mice where both tumors reached 250-300 mm^3^ in volume (measured by 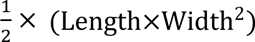 × (Length×Width^2^)) were selected for further experimentation. For tumor growth monitoring, mice were treated with vehicle, CV, or ASP9801 at D0, D2, and D4. αPD-1 (Bioxcell, #BE0146) or IgG isotype control (Bioxcell, #BE0089) was administered weekly at 200 µg/mouse via IP injection. Tumors were measured every other day under anesthesia and with a calibrated digital caliper until tumor volumes reached a cumulative tumor burden of approximately 3000 mm^3^ or humane endpoint. Mice that succumbed to potential contamination/microbial infection at tumor site were excluded from analysis. For single-cell RNA sequencing and flow cytometry, mice were treated with 30 μL PBS (vehicle) intratumor (IT) in the right-sided tumor, 2 x 10^7^ PFU in 30 µL of CV (IT) in the right-sided tumor and 200 µg of Isotype control (IP injection), 30 µL of PBS (IT) in the right-sided tumor and 200 µg of αPD-1 (Anti-PD-1 Clone, RMP1-14) (IP injection), 2 x 10^7^ PFU in 30 µL of ASP9801 (IT) in the right-sided tumor and 200 µg of isotype control (IP injection), 2 x 10^7^ PFU in 30 µL of ASP9801 (IT) in the right-sided tumor and 200 µg of αPD-1 (IP injection). At day 7, and tumors and blood were harvested for sample preparation.

### PDX generation for tumor slice experiments

NSG (NOD.Cg-*Prkdc^scid^ Il2rg^tm1Wjl^*/SzJ) (The Jackson Laboratory, #005557) mice were housed, handled and bred in in accordance with protocols approved by the Houston Methodist IACUC committee. These mice were injected subcutaneously with PDX suspension on their flanks using a 0.5 mL syringe equipped with a 26-gauge needle. To prepare the tumor suspension, cryopreserved PDX fragments were quick thawed in a 37C° incubator, washed with DPBS to remove traces of DMSO, and minced with a razor blade to generate a slurry with tumor pieces that would be able to be drawn into the syringe. This suspension was transferred to a 1.5 mL Eppendorf tube and resuspended with enough DPBS so that each mouse could be injected with 100 μL of tumor suspension. Injected mice were monitored daily for tumor growth and harvested once tumors reached a 500 - 1000 mm^3^ in volume. For a list of PDX models used, see **Supplementary Table 2**.

### Stereotaxic intracranial injections GBM PDX model generation

NSG were purchased from the JAX repository or bred in-house. Mice were housed and handled in accordance with protocols approved by the Houston Methodist IACUC committee. For intracranial injections, 6-8 weeks old NSG mice were used. Briefly, single cell suspensions were prepared by digesting patient tumor or PDX fragments with Accutase (VWR, #10210-214) at 37C° for 10-15 minutes. Tumor digests were washed with D-PBS to remove Accutase and resuspended at a concentration of 1×10^5^ per µL, and 1 µL was injected into the striatum of each mouse using a NeuroStar’s Drill and Injection stereotaxic device (Bregma: −0.5/2.5/-3.5). Mice were harvested once they began showing signs of neurological decline such as ataxia, severe lethargy, tilted heads, hunched posture.

### Ex vivo tumor slice culture and treatment

Tumor slice assays were performed with PDX and patient tumor samples according to a proprietary protocol licensed to EMPIRI, inc. Treatment using Astellas virus was performed by adding 10,000 PFU of virus directly onto E-slices. Viability changes were measured on day 1 (baseline) and on days 8 and 12 in culture using the E-slice method. E-slice assay culture method is proprietary but available for researchers through EMPIRI, inc.

### In vitro cell-line response to Astellas virus

MC38, CT26, U87, Detroit562, A549, and HCT116 cell lines (see **Supplementary Table 1**) were cultured in 10 cm plates in DMEM (GenDEPOT, #CM002-050) or RPMI-1640 (GenDEPOT, #CM059-050) supplemented with 10% FBS (GenDEPOT, #F0601-050) and passaged for use at ∼50% confluence. These were seeded at 10,000 cells per well in 100 μL of media in sterile treated 96-well plates. After allowing the cells to settle and adhere, viability readings were taken at 4, 24, 48 and 72 hours after seeding. Virus treatment was added after the 4-hour viability reading at a concentration of 0.01 and 0.05 MOI. Assessing viability was performed with CellTiter-Blue® reagent (Promega, G8081-100ml) diluted 1:10, of which 5 μL were added to each well for a final dilution of 1:200. For 96- well plate assays, the CellTiter-Blue® reagent was left to incubate at 37C for 4 hours before reading. To not disturb the adherent cells with repeat viability readings, separate triplicates were prepared for each time point. Wells with signs of media evaporation were excluded or uneven CellTiter-Blue administration were excluded from analysis.

### Histology analysis

Tissues were fixed in 10% formalin (Fisher Scientific, #11-002-205) then washed with 1 mL 70% ethanol before being placed in cassettes and embedded in paraffin blocks by the HMRI Pathology Core. The pathology core also processed the provided samples by sectioning them into 5 μm sections and stained with hematoxylin and eosin for histological analysis. Alternatively, some samples were fixed in 4% paraformaldehyde overnight and embedded in OCT for frozen sections (10µM).

### Immunofluorescence analysis

Anti-vaccinia virus antibody (Abcam, #Abs35219) was used in standard immunofluorescence analysis on 10µM thick frozen sections from E-slices, using a secondary antibody conjugated with Alexa Fluor 488 (Fisher Scientific, #A-11017). Nuclei were visualized by counterstaining with DAPI. DsRed fluorescence is from the transgene.

### Single-cell RNA sequencing analysis

Single-cell RNA libraries were generated from dissociated tumors following the 10x Genomics Chromium Next GEM Single Cell 5’ Reagent Kits v2 (Dual Index) protocol (RRID:SCR_022645). Raw Illumina sequencing reads were aligned to the mm10 (mouse) reference genome using the Cell Ranger pipeline (v9.0.0, RRID:SCR_017344) with default parameters. Downstream analysis was performed using the filtered feature-barcode matrices. To ensure high-quality data, low-quality cells were removed via miQC (v1.1.0, RRID:SCR_022697, “https://github.com/greenelab/miQC”) using a posterior probability cutoff of 0.8. Potential doublets were identified and excluded using scDblFinder (v1.16.0, RRID:SCR_022700), with the doublet formation rate adjusted according to the total number of cells recovered in each library. Following QC, data were normalized, integrated, and clustered using Seurat (v5.2.1, RRID:SCR_016341). To account for batch effects across treatment sites and experimental runs, integration was performed using Harmony (v0.1.0, RRID:SCR_022206). Gene expression counts were normalized to library size and log2-transformed. Dimensionality reduction was performed via Principal Component Analysis (PCA) using the top 2,000 highly variable genes. Unsupervised clustering was conducted using a Louvain-based graph approach on the Harmony-corrected principal components, with resolution parameters optimized between 0.1 and 1.0 for each dataset, cluster-specific marker genes were identified using a fast Wilcoxon AUC rank-sum test (presto R package v1.0.0 “github.com/immunogenomics/presto”) and visualized through dot and feature plots. Functional enrichment analysis was performed to characterize the biological pathways associated with differentially expressed genes (DEGs). Gene Ontology Biological Process (GO_BP) and MSigDB (RRID:SCR_016863) Hallmark gene set analyses were conducted using the SCP R package (v0.5.6, github.com/zhanghao-njmu/SCP). Cellular identities were assigned through a combination of marker gene expression analysis and automated annotation using SingleR (v2.4.1, “github.com/dviraran/SingleR”) against the ImmGen reference database, as well as reference-based projection via ProjecTILs (v3.3.0, “github.com/carmonalab/ProjecTILs”) for the T-cell compartment. To distinguish malignant cells from the tumor microenvironment, we performed large-scale chromosomal copy number variation (CNV) analysis using inferCNV (v1.18.1, “github.com/broadinstitute/infercnv”). Non-malignant immune cells were utilized as a reference to define the baseline genomic state.

### TCR Analysis with Immunarch and scRepertoire

T-cell receptor (TCR) sequences were analyzed by integrating Cell Ranger V(D)J output files with transcriptomic data. Filtered contig data were processed and visualized using the Immunarch R package (v0.9.1, RRID:SCR_023089). For the integration of TCR clonotypes with single-cell gene expression profiles, the scRepertoire R package (v2.2.1, github.com/BorchLab/scRepertoire) was utilized. Clonotypes were defined based on the combined amino acid sequences of both the alpha and beta chains. Integration with the Seurat object was performed as outlined in the scRepertoire documentation without modification to the default parameters.

### Flow Cytometry

Freshly dissected mouse tumor tissues were briefly washed in PBS, minced into small 1- 2 mm pieces, and treated with an enzyme digestion mixture of 20% Collagenase/Hyaluronidase (GenDEPOT, CA094-002) and 100 μg/mL DNAse I in Accutase for shaking 800 RPM for 30 minutes at 37°C. After centrifuging at 600 RCF, the enzyme mix was removed, digested tissues were resuspended in DMEM/F12+B27 to generate single cell suspensions and filtered through a 40 μM cell strainer (ThermoFisher Scientific, #NC1789427). Cell suspensions were then resuspended in RBC lysis buffer for 10 minutes on ice to remove red blood cells. Cells were then resuspended in DMEM/F12+B27 (ThermoFisher Scientific, #17504044) and layered on 5 mLs of PBS buffered 20% Percoll (Sigma Aldrich, #P4937-500ML), centrifuged at 800 RCF and 4 C for 30 to remove cell debris. The cell pellet was then washed, resuspended in DMEM/F12+B27, and counted. 1 million cells/sample was taken and resuspended in brilliant stain buffer containing 1:100 dilution of mouse TruStain FcX (Biolegend, 101320). After 10 minutes of blocking at 4°C, cells were stained with flow cytometry antibody cocktails for 40 minutes, washed and stained with Zombie Aqua (Biolegend, 423104) for 20 minutes, fixed with IC fixation buffer (ThermoFisher Scientific, 00-8222-49), and analyzed using a BDFacs Symphony A5SE. Some MC38 tumor samples were excluded from analysis due to poor quality or low amount of tumor present at harvest (low number of viable cells from tumor digestion). Spectral flow cytometry data was analyzed in Flowjo (v10.10.0) and plotted in Graphpad Prism (v11.0.2).

| Fluorophore-Antibody | Catalog number | Dilution |
| --- | --- | --- |
| AF488-CD25 | 102017, Biolegend | 1:1000 |
| APC-CD3 | 156106, Biolegend | 1:1000 |
| APC-Fire750-MHC Class II (I-A/I-E) | 107652, Biolegend | 1:1000 |
| APC-Fire810-Ly6c | 128055, Biolegend | 1:1000 |
| BUV495-NK1.1 | 364-5941-82, eBioscience | 1:500 |
| BUV615-CD11c | 366-0114-82, eBioscience | 1:500 |
| BUV661-CD8 | 569186, BD | 1:500 |
| BUV805-CD45 | 3680452-82, eBioscience | 1:1000 |
| BV421-CD86 | 105032, Biolegend | 1:500 |
| BV510-Zombie Aqua | 423101, Biolegend | 1:1000 |
| BV605-PD-L1 (CD274) | 124321, Biolegend | 1:500 |
| BV650-CD69 | 104541, Biolegend | 1:500 |
| BV750-CD4 | 100467, Biolegend | 1:1000 |
| BV785-F4/80 | 123141, Biolegend | 1:500 |
| PE -TIM-3 (CD366) | 134003, Biolegend | 1:500 |
| PE-Cy7 -PD-1 (CD279) | 109109, Biolegend | 1:1000 |
| PE-Dazzle-CD11b | 101255, Biolegend | 1:1000 |
| PE-Fire 810-Ly-6G | 127673, Biolegend | 1:1000 |
| PerCP-Cy5.5-CD44 | 103031, Biolegend | 1:1000 |
| Spark UV 387-CD19 | 115585, Biolegend | 1:1000 |

### Cytokine analysis from tumor slice conditioned media

Conditioned media collected at day 8 or 12 from E-slices were maintained at −80C until time for analysis. These tubes were shipped to RayBiotech for quantitation using the Human Cytokine Array Q640 multiplex immunoassay (Raytech, #QAH-CAA-640).

Absolute amounts of secreted proteins were measured and recorded as concentration values. For correlation analysis, cytokine concentration values were paired with percent change in viability of treated sample compared to the untreated sample. Correlation was assessed using the Pearson product-moment correlation test from R package ggpubr (v0.6.0). For differential secretion analysis, cytokines were analyzed both by comparing concentrations directly and through the R package Limma (v3.62.2) using a custom pipeline. For Limma analysis, cytokine values were imported along with corresponding clinical metadata and treatment response status. Cytokines whose expression was undetectable in more than half the samples were filtered out. Cytokine concentrations were then log normalized. Samples were then split by responder vs non-responder status as determined by cell viability experiment results. Normalized expression values were fitted to a linear model based, and empirical Bayes moderation was applied to compute moderated t-statistics and log-fold changes. Adjusted p values were generated for multiple testing using the Benjamini-Hochberg method. For direct comparison of cytokine concentrations, we split samples by responder vs non-responder status and assessed differences using a student’s t test. To assess pairwise changes in cytokine secretion between baseline and ASP9801-treated samples, we subtracted cytokine concentrations of ASP9801-treated samples from baseline samples in a pairwise manner, split by samples by responder vs non-responder status, and differences were assessed using a Welch’s two sample t-test. For principal component analysis (PCA), we used the R package PCAtools (v2.18.0). Cytokines concentration values were again log normalized. PCA was performed using singular value decomposition as implemented in the PCAtools pca() function. Sample-level metadata were incorporated to enable downstream visualization and annotation.

### Statistical analyses

Statistical comparisons were performed using GraphPad Prism (v11.0.2) or R. Values and error bars represent the mean ± standard error of the mean, unless otherwise stated. For violin plots, ends represent minimum and maximum values, solid line represents median value, and dashed lines represents quartiles. *p*-values were determined by an appropriate statistical test such as student’s *t*-test or ANOVA with multiple comparison correction, as indicated in figure legends.

### Data Availability

All data are included in the Supplementary Information or available from the authors; source data for charts and graphs is provided with this paper. Single-cell RNA sequencing data generated in this study will be deposited at GEO and made publicly available upon publication.

## Acknowledgements

We thank Geoffrey Hummelke for editing this manuscript.

## Supplementary Figures

**Supplementary Figure 1.**
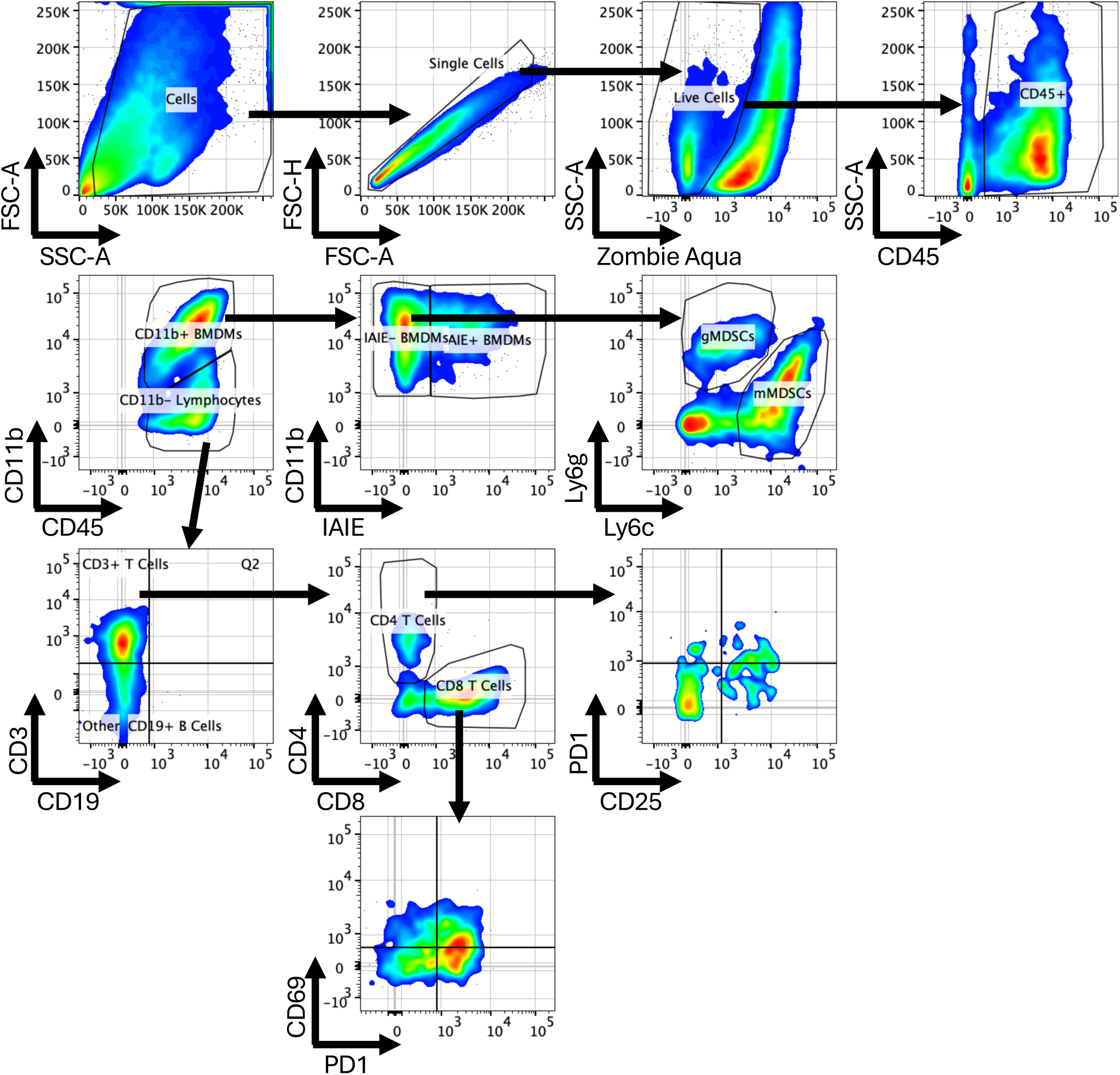
Gating strategy used for analyzing intratumor immune cells, performed using Flowjo software (v10.10.0).

**Supplementary Figure 2.**
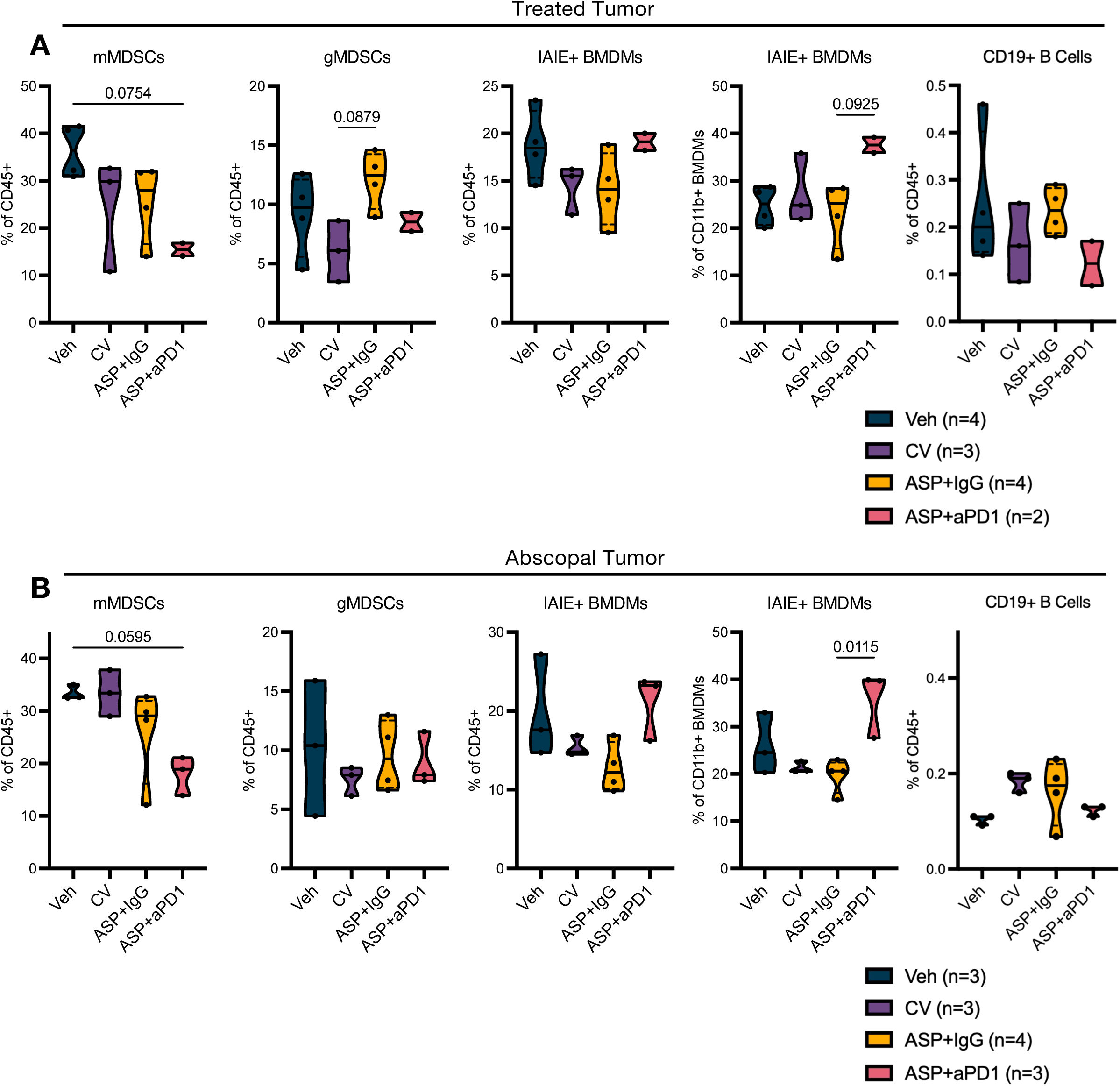
Flow cytometry analyses of treated and abscopal MC38 tumors. **A-B,** Flow cytometry analysis of intratumor myeloid cells from treated tumor (A) and abscopal tumor (B). For violin plots, ends represent minimum and maximum values, solid line represents median value, and dashed lines represents quartiles. *p*-values determined by one-way ANOVA with Šidák’s correction for multiple comparisons.

**Supplementary Figure 3.**
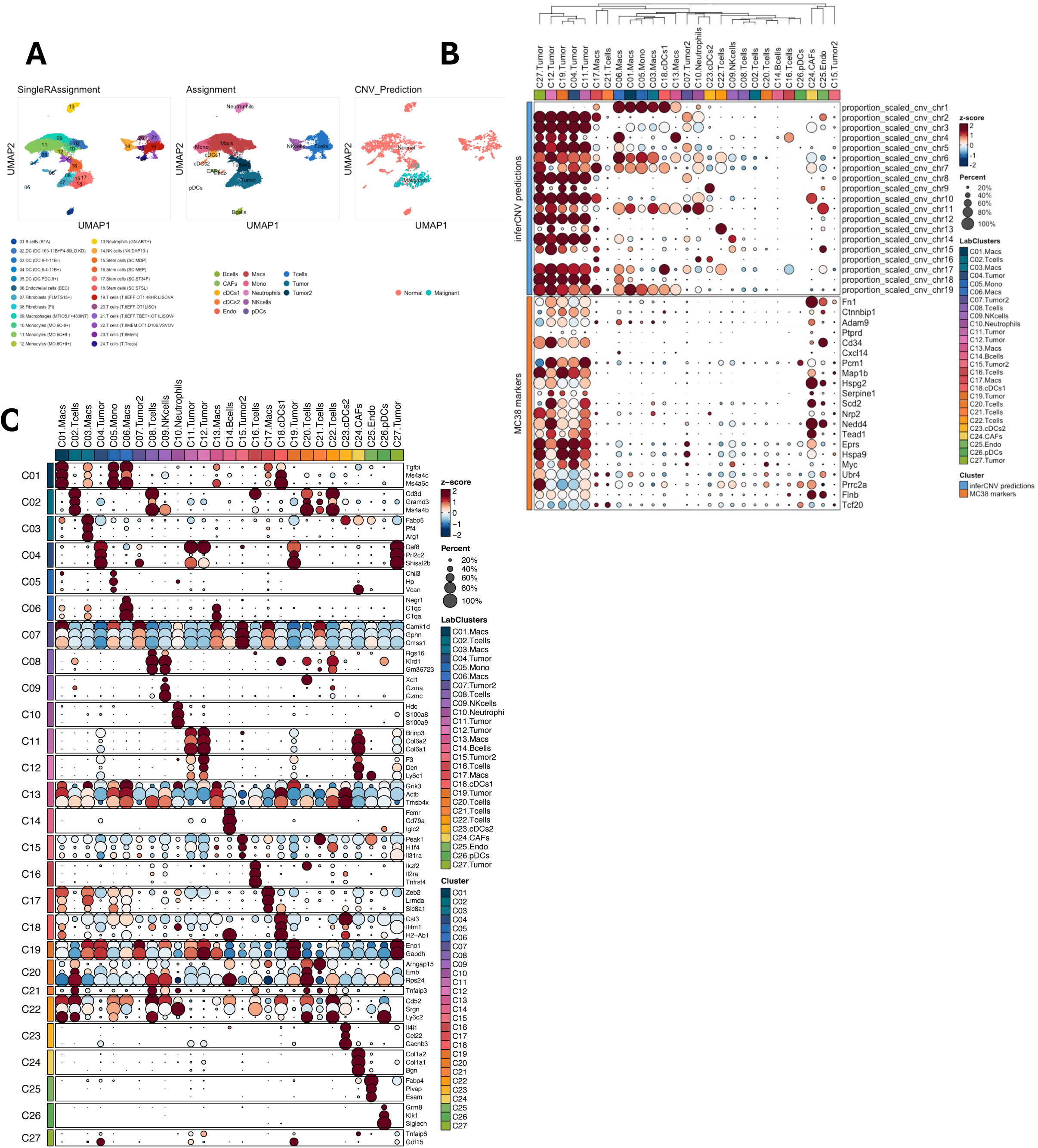
Single-cell transcriptomic profiling and malignant cell identification. **A,** UMAP projections of 40,871 integrated single cells, annotated by SingleR, manual lineage assignment, and InferCNV-predicted malignancy status. **B,** Dot plot showing InferCNV-estimated proportions alongside MC38-specific marker gene expression. **C,** Expression profiles of top differentially expressed genes (DEGs) by cluster. Dot size represents the percentage of cells expressing the gene; color intensity indicates the Z-score of expression or InferCNV-estimated proportions.

**Supplementary Figure 4.**
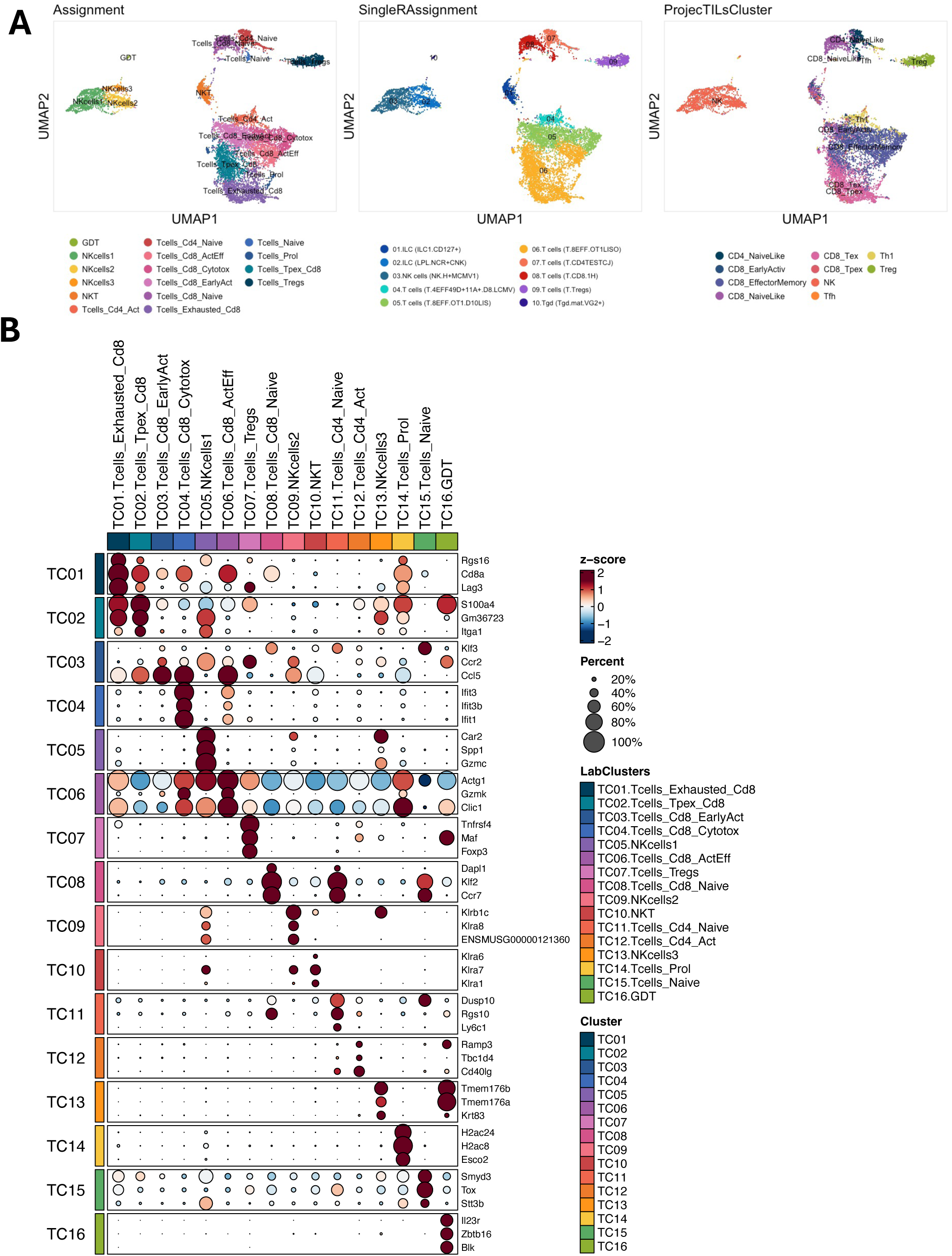
Single-cell transcriptomic profiling of T and NK cells. **A,** UMAP visualization of T and NK cell sub-clusters (n = 10,495) colored by manual, SingleR, and ProjecTILs cell-type assignments. **B,** Dot plot of top DEGs characterizing the T and NK cell compartments. Dot size represents the percentage of cells expressing the gene; color intensity indicates the Z-score of expression.

**Supplementary Figure 5.**
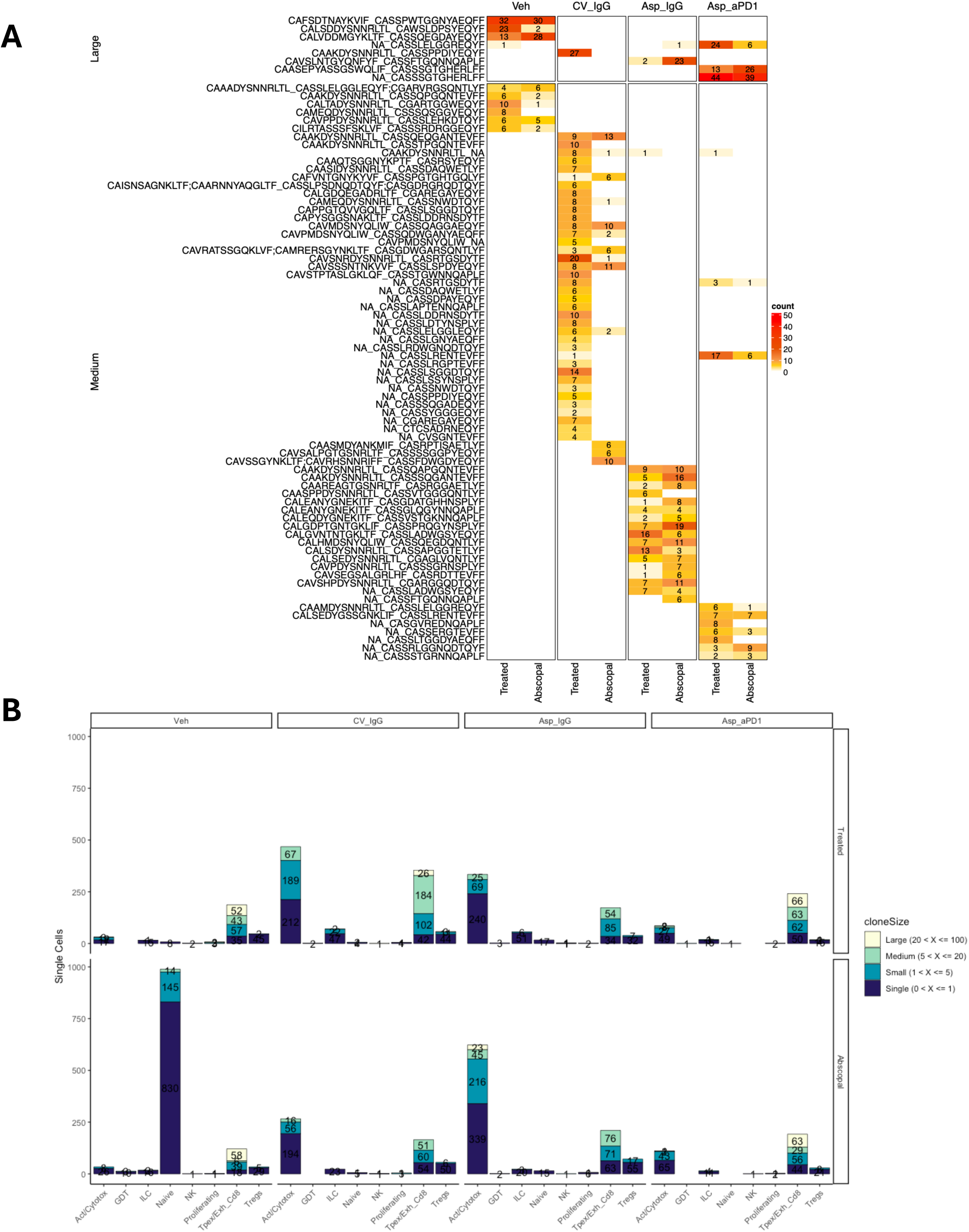
Single-cell transcriptomic profiling of intratumor T cell clones. **A,** Heatmap showing the count of large and medium CTaa clones per Sample. **B,** Stacked bar plots showing the number of cells per cell type and colored by cloneSize.

**Supplementary Figure 6.**
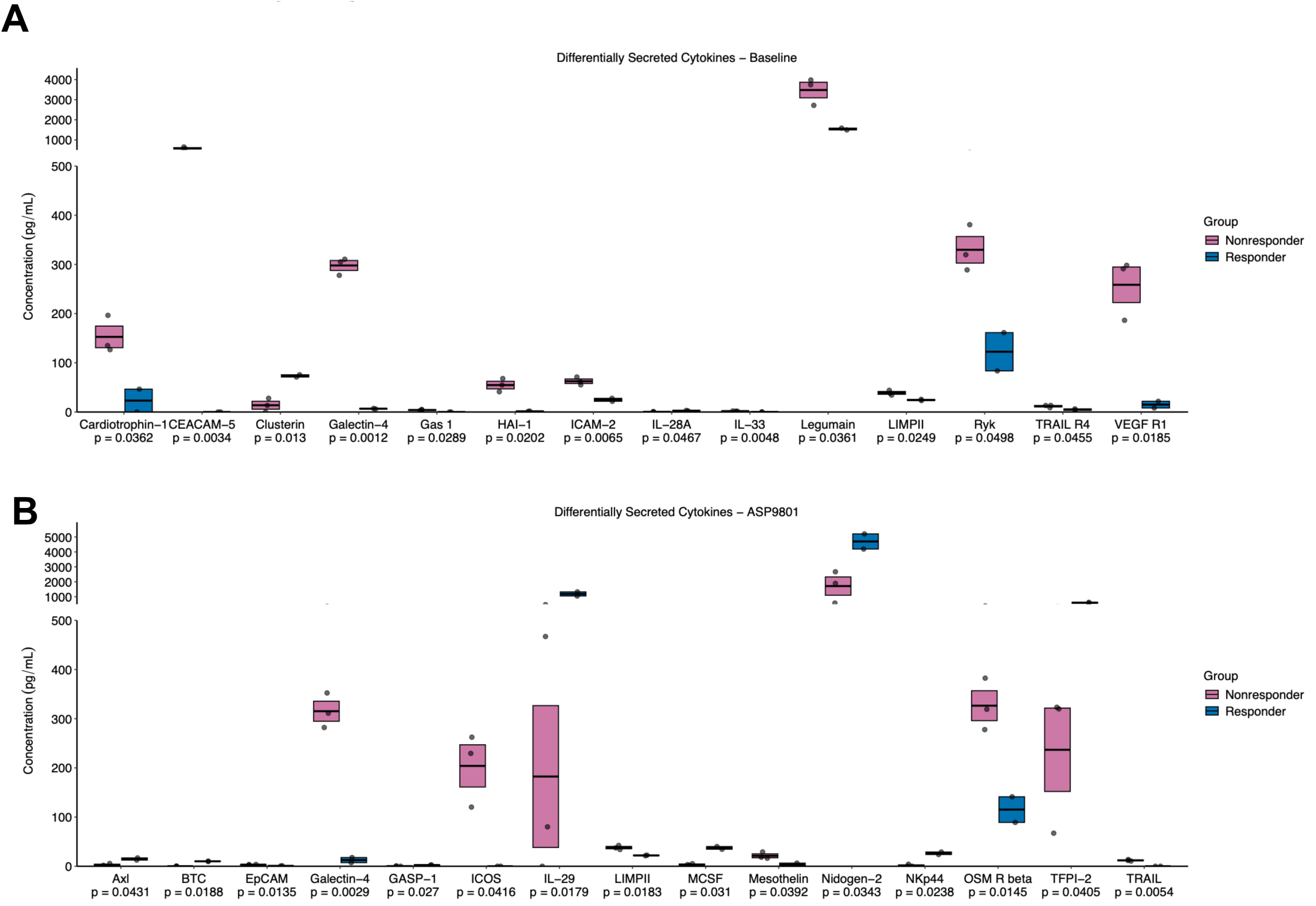
Cytokine Analysis for Virus Treated Experiments. **A-B** Cytokine array quantitation of tumor slice conditioned media for differentially secreted cytokines between non-responders (n=3) vs. responders (n=2) by direct comparison, presented as pg/ml. (A) patient samples at baseline (B) ASP9801 treated patient samples. Floating box represents mean with SEM, with horizontal line indicating the mean; individual values are overlaid as points. *p*-values determined by two-sample unpaired *t*-test.

